# Strong trait correlation and phylogenetic signal in North American ground beetle (Carabidae) morphology

**DOI:** 10.1101/2021.02.12.431029

**Authors:** Jacob D. Stachewicz, Nicholas M. Fountain-Jones, Austin Koontz, Hillary Woolf, William D. Pearse, Amanda S. Gallinat

## Abstract

Functional traits mediate species’ responses to and roles within their environment, and are constrained by evolutionary history. While we have a strong understanding of trait evolution for macro-taxa such as birds and mammals, our understanding of invertebrates is comparatively limited. Here we address this gap in North American beetles with a sample of ground beetles (Carabidae), leveraging a large-scale collection and digitization effort by the National Ecological Observatory Network (NEON). For 154 ground beetle species, we measured seven morphological traits, which we placed into a recently-developed effect-response framework that characterizes traits by how they predict species’ effects on their ecosystems or responses to environmental stressors. We then used cytochrome oxidase one sequences from the same specimens to generate a phylogeny and tested evolutionary tempo and mode of the traits. We found strong phylogenetic signal in, and correlations among, morphological ground beetle traits. These results indicate that, for these species, beetle body shape trait evolution is constrained, and phylogenetic inertia is a stronger driver of beetle traits than (recent) environmental responses. Strong correlations among effect and response traits suggest that future environmental drivers are likely to affect both ecological composition and functioning in these beetles.

## Introduction

Functional traits mediate interactions between organisms and their environment (Violle et al. 2007), and therefore play a key role in our understanding of ecosystem functions, services, and environmental responses (McGill et al. 2006, Díaz et al. 2013). The general patterns of functional trait macro-evolution are remarkably variable, showing both strong evolutionary conservatism and lability, with consequences for subsequent ecological assembly (Webb et al. 2002, Ives and Godfray 2006, Cavender-Bares et al. 2009). For example, ecological selection on strongly conserved traits (where species resemble their close relatives (Pagel 1999, Wiens et al. 2010) naturally leads to ecological communities with reduced phylogenetic diversity. Such low-diversity communities are typically less productive and stable (but not always; see Tucker et al. 2017). The converse may be true for evolutionarily labile traits, particularly clades that have undergone adaptive radiations (Seehausen 2006) or convergence (Muschick et al. 2012). Yet it is rare for macro-evolutionary studies to contrast the evolutionary tempo and mode among traits driving ecological assembly and function (but see Díaz et al. 2013), and rarer still for non-plant taxa.

The effect-response framework has emerged within ecology as a powerful, if somewhat controversial (Savage et al. 2007, Luck et al. 2012), paradigm for understanding how the traits driving species’ ecological assembly might be linked with their ecological functions (Díaz and Cabido 2001, Suding et al. 2008). Species’ effect traits are associated with their ecological impact, including resource use, habitat modification, and contributions to nutrient cycling. Response traits, on the other hand, mediate how species are affected by their environments, including dispersal, colonization, and persistence within and across sites. They can be used to make predictions about species’ responses to environmental stressors and potential for re-invasion following extirpation. The effect-response framework has thus gained traction in ecological risk-assessment for predicting the consequences of environmental change for (plant and arthropod) species (Díaz and Cabido 2001, Wong et al. 2019). In such risk-assessment, strong correlations among effect and response traits are of concern, because they imply that species lost due to environmental change (on the basis of response traits) will impact ecosystem function (via effect traits). Beyond current ecological uses, the evolutionary history of response traits can provide an informative context for how species and their traits respond across environmental gradients (Poff et al. 2006) Accordingly, effect-response trait-based risk assessment has been extended to consider phylogenetic patterns, both to predict species’ effect and response traits, and also to highlight where conserved trait evolution would suggest higher risk through correlation among traits (Díaz et al. 2013).

The phylogenetic extension of the effect-response framework has been primarily applied to plants and large vertebrates, leaving a conspicuous gap in our understanding of phylogenetic patterns of effect and response traits for invertebrates. This mirrors a broader limitation in our understanding of macro-evolutionary processes across systems due to taxonomic biases toward clades such as mammals, reptiles, and trees (Harmon et al. 2003, Harmon et al. 2010, Eastman et al. 2011). Insects, particularly ground beetles (Carabidae), offer many species-rich and ecologically important clades for investigating scale-dependence in phylogenetic patterns. Clade-specific studies of phylogenetic signal in beetle functional traits have revealed a variety of patterns, such as variation in phylogenetic structure with trophic position, with predators being more constrained than detritivores (Fountain Jones et al. 2017), and strong phylogenetic signal in body size-structured communities of diving beetles (Vamosi and Vamosi 2007). And while beetle lineage diversity appears to have been generated early in their history (Hunt et al. 2007), it is not clear whether this radiation of lineages matches a radiation of traits. Previous studies that have focused on beetle clades have revealed the interplay between the evolution of species’ traits and their present-day ecology. This has included evolutionary conservatism of environmental tolerance (Hortal et al. 2011), habitat adaptations (Vamosi and Vamosi 2007), and environmentally filtered traits (Gossner et al. 2013). Understanding the role of beetle functional trait evolution in species assembly and ecosystem function is further warranted by the ecological importance of beetles as biological links between producers and consumers (Kotze et al. 2011), ecosystem engineers, (Logan and Powell 2001, Albert et al. 2012) and high-impact pests (Bentz et al. 2010). Ground beetles (Carabidae) in particular are frequently used as indicator species due to their rapid responses to habitat fragmentation and degradation (Ribera et al. 2001, Rainio and Niemelä 2003), local-scale attraction to high-quality resources (Haila et al. 1994), and economically significant predation of agricultural pests (Hance et al. 1990, Collins et al. 2002). Exploring the links between beetles’ response and effect traits would generate insight into ecosystem function under environmental change; incorporating phylogeny may allow prediction for the thousands of species about which we know comparatively little (Cardoso et al. 2011). With recent work explicitly placing beetle traits within the effect-response framework (Fountain Jones et al. 2015, Moretti et al. 2017), it is both timely and worthwhile that we apply this framework to understanding the evolutionary history and ecological consequences of beetle traits across North America.

We applied the effect-response trait framework to test the macro-evolution of functional traits for North American ground beetles. We quantified seven morphological traits from 1975 beetle specimens collected by the National Ecological Observation Network (NEON), and built a dated phylogeny from 2399 DNA sequences, using molecular data from the measured (and additional) specimens. For 154 species, we quantified the tempo and mode of beetle trait macro-evolution and tested whether trait evolution maps onto ecological function within the effect-response framework. The results of this analysis are intended to inform ecological risk assessment for North American beetles, and to provide an open-access dataset for further exploration of beetles in the NEON field system.

## Materials and Methods

All data collected as part of this study are released in the Supplementary Materials, along with R code to reproduce analyses and figures (Appendix S1).

### Effect-response framework and measurement

We selected traits for this study based on an existing literature review that characterizes well-studied beetle traits into an effect-response framework (Fountain Jones et al. 2015). We note that how traits affect ecosystem services is poorly understood for beetles, and arthropods in general (Fountain-Jones et al. 2015, Wong et al. 2019). We chose seven morphological traits that were possible to accurately assess from images of specimens (see below) and that were previously classified as effect and/or response traits (Table 1).

**Table 1.**
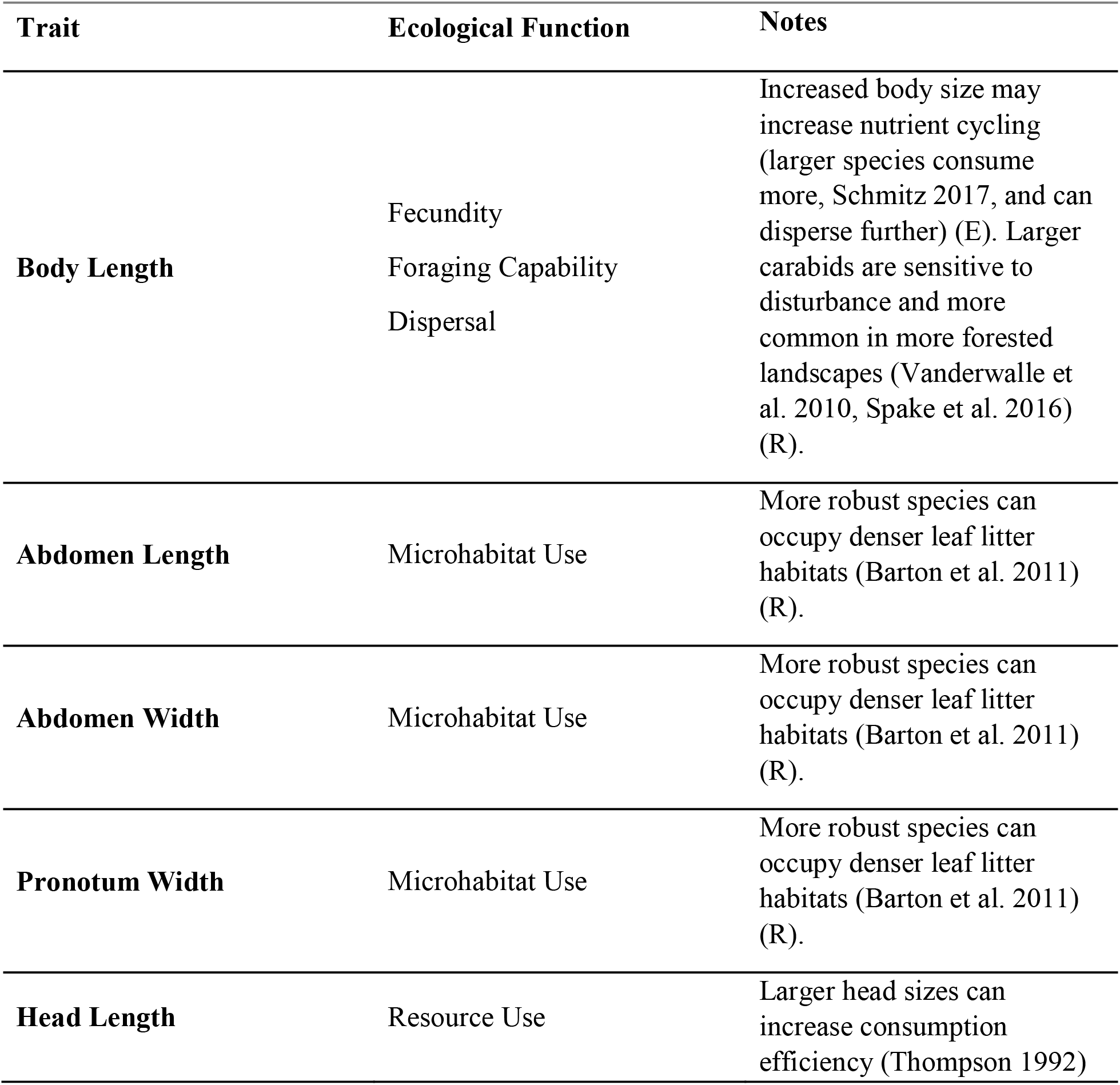
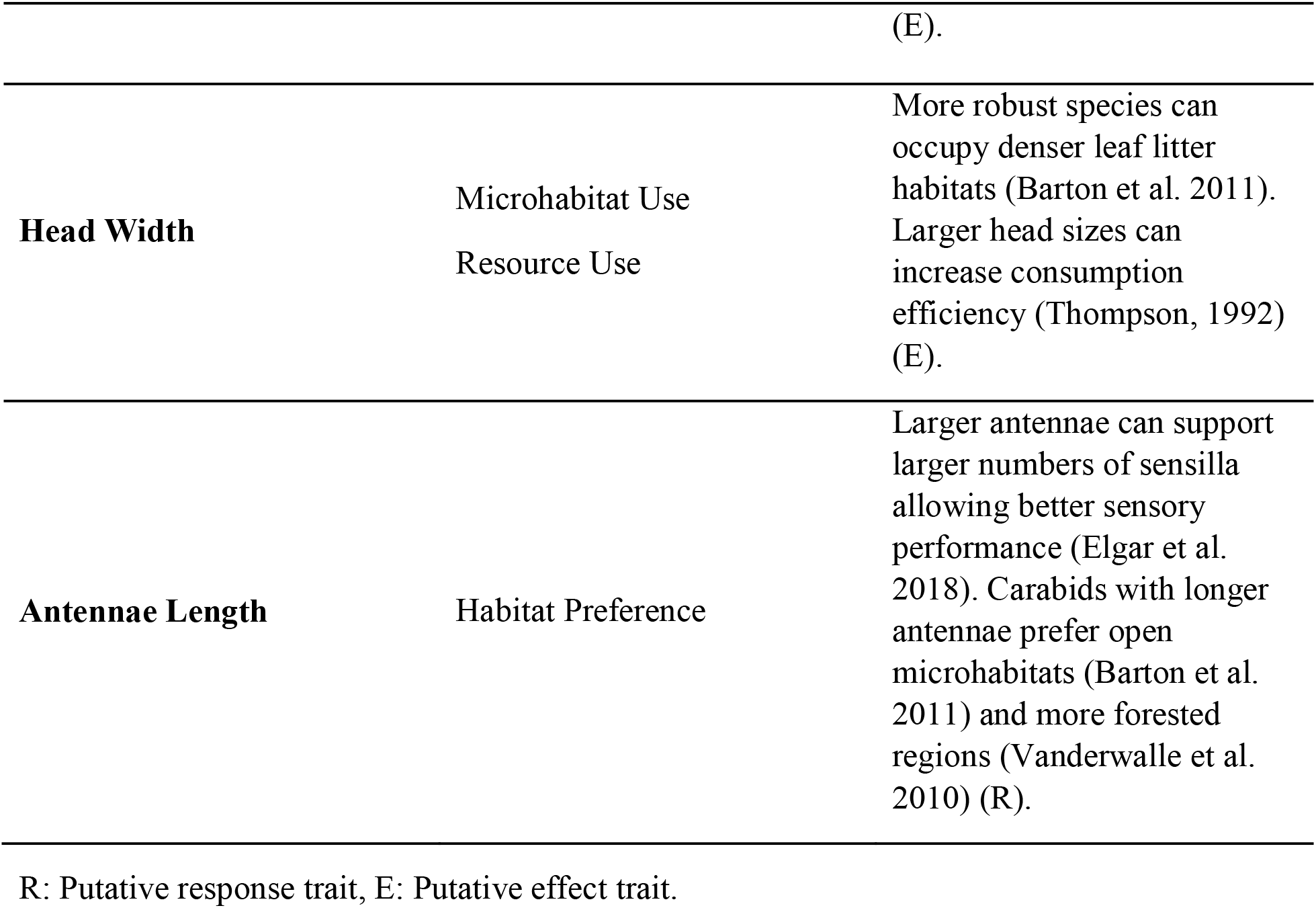
Classification of seven morphological ground beetle traits in the effect-response trait framework. For each trait we identify an associated ecological function, and whether the trait is linked to a species’ *effect on* (E) or *response to* (R) its environment, based on classifications from Fountain□Jones et al. (2015). Different traits can map onto the same ecological function, and a trait can be both an effect and response trait.

We measured the seven target morphological traits using dorsal images of ground beetle specimens (one image per specimen) provided by NEON. The specimens were sampled using pitfall traps across NEON’s terrestrial North American field sites, which are spread geographically from Alaska to Puerto Rico and represent a wide range of environmental conditions (Appendix S2). We measured traits using the program FIJI (Schindelin et al. 2012), standardizing the images with a 5 mm scale to account for variation among photos, and recording all traits at the maximum possible length or width for each feature. All 13,825 observations from 1975 specimens are available in the Supplementary Materials (Appendix S3). Species identifications were performed by NEON taxonomists. For each species, we calculated median trait values (mean = 12 individuals per species) for use in subsequent analysis. Information regarding beetle sex was not recorded. We included 154 species in all trait analyses, and we note that 12 of what we refer to as “species” throughout are in fact morphospecies, often only identified to the genus level. We have included those groups in our analyses because sequences provided from the specimens allowed us to place them in the phylogeny.

All trait measurements were made by a single observer, using a protocol generated as part of this study (Appendix S4). A major advantage of our dataset is that each specimen was identified and DNA sequenced (see below), giving a high degree of taxonomic precision and reproducibility to our analysis. Our protocol was, however, designed around the limitation that the only images we have of the beetles are dorsal views, which imposes limitations on the measurements we were able to make. While these measurements are, therefore, limited, we feel the comprehensive coverage of this dataset in terms of geography, taxonomy, and genetic data give us a unique opportunity to examine the morphology of these beetles.

### Phylogeny construction

NEON have released cytochrome oxidase one (COX1, the barcoding region for beetles) sequences for all of the specimens digitized and measured in this study as well as some additional specimens (Hebert et al. 2003). We used these sequences to generate a phylogeny of our species (along with five outgroup sequences from *Myrmeleon immaculatus*, *M. formicarius*, *Brachynemurus abdominalis*, *B. ferox*, and *Dendroleon obsoletus*). We aligned these sequences using MAFFT (Katoh 2002), and then performed 1000 independent maximum likelihood searches using RAxML under a GTR-GAMMA model (using a re-arrangement ‘i’ parameter of 10, chosen through likelihood comparisons as outlined in the RAxML manual; Stamatakis 2014; Appendix S6). The most-likely tree was then dated using treePL (Smith and O’Meara 2012), with the smoothing parameter estimated using cross-validation, using a date range of 290-310 mya for the outgroups (Zhang et al. 2018). This single, dated phylogeny (Appendix S7) was then used for all downstream analysis. We emphasize that this phylogeny, which was built using a single locus, should not be viewed as a definitive phylogeny for North American beetles; while we have excellent taxonomic coverage (i.e., species collected from across North America) that is not matched by genetic coverage (i.e., a single barcoding region). Our goal with this phylogeny (which is, itself, a hypothesis) is to use it as a tool to test a series of questions about the tempo and mode of ground-beetle morphological evolution.

### Modeling the evolution of individual traits

The last few years have seen an explosion in so-called ‘phylogenetic comparative methods’, which can be used to test hypotheses and questions about the tempo (speed) and mode (kind) of trait evolution (reviewed in Cooper et al. 2010). While our ultimate goal is to understand how beetle morphology may have co-evolved, and to use the effect-response framework to consider the consequences of such co-evolution on ecological function, we begin by verifying our underlying data using classic comparative methods. We therefore begin by fitting explicit statistical models to test the most likely kind (mode) of trait evolution, using information theoretic criteria (AIC) to assess models’ fit while accounting for their different degrees of complexity (Boettiger et al. 2012). Our first model is a ‘white noise’ (null) model that tests whether there is any association between each trait and our estimated phylogeny; since beetle morphology has been used for centuries to assign taxonomy (and that taxonomy broadly maps onto phylogeny), strong support for this model would suggest our phylogeny is inaccurate. Thus this first model test is critical to verifying our approach. We then estimated the fit of Pagel’s λ and δ (Pagel 1999) to these data; λ measures the extent to which the data are consistent with a Brownian Motion (BM) model of evolution (a λ of 0 indicating no fit, and 1 indicating pure BM), and δ the extent to which the data fit a model of accelerating or decelerating evolution (δ<1 indicating decelerating, and δ>1 indicating accelerating). These transformations are notable for being relatively sensitive to the presence of any phylogenetic signal (the tendency for closely-related species to resemble one-another) and to be able to assess the magnitude of that resemblance (Münkemüller et al. 2012). We fit pure models of Brownian Motion evolution (consistent with inherited, non-directional trait evolution and comparable with Pagel’s λ; see Revell et al. 2008 for further discussion), trends in increasing or decreasing rates of evolution through time (to be compared with Pagel’s δ), which represent the special cases of Pagel’s λ and Pagel’s δ, respectively, where there is perfect signal. Finally, we fit a single-optimum Ornstein-Uhlenbeck model (Butler and King 2004), which is broadly thought to test for clade-level trait optima (Uyeda and Harmon 2014).

All models were fit using *geiger* (Harmon et al. 2008).

### Modeling the co-evolution of traits

Species traits rarely, if ever, evolve independently, and to test for broad patterns in, and the evolution of, beetle allometric relationships we conducted a phylogenetic Principal Components Analysis (Revell 2009). A standard PCA identifies the major axes of variation within a dataset of continuous traits, identifying how multiple variables (in this case traits) are correlated with one-another. So, for example, if beetles with longer legs tended also to have longer and narrower carapaces, then a PCA might detect and represent this pattern by having an axis with strong positive loadings (essentially correlations) with leg and carapace length, and a negative loading with carapace width. A phylogenetic PCA is analogous, but differs in accounting for how species’ shared evolutionary history makes them non-independent observations (Cooper et al. 2010). This allows us to account for and measure the degree to which observed associations are likely the result of shared, inherited trait variation. The resulting principal component axes can be analysed analogously to ‘standard’ PCA axes, but we additionally have a measure of phylogenetic signal – Pagel’s λ – that is interpreted exactly as it is when applied to univariate data (see above).

## Results

Among the 154 species tested from 50 genera, the traits with the greatest variation were body length (SD = 5.60, ranging from *Elaphropus anceps,* 2.43 mm, to *Pasimachus strenuus,* 28.16 mm) and abdomen length (SD = 3.43). The trait that varied the least among species was head length (SD = 1.37). We found strong evidence of phylogenetic signal (all λ > 0.73; all p < 0.001) in all traits measured (Table 2), providing supporting evidence that our phylogeny (despite the limitations outlined in the methods) is sufficient to detect pattern and process in trait macro-evolution. λ models best explained beetle body shape trait evolution based on AIC scores for all seven traits (Table 2). Strong support for Pagel’s λ is consistent with trait evolution intermediate between pure chance and pure Brownian motion, *i.e.*, phylogenetic signal. Phylogenetic PCA further revealed that all measured body shape traits are highly correlated regardless of classification as an effect or response trait (Figure 1; Appendix S8). PC1 accounted for 86.1 percent of the component variance, while PC2 accounted for an additional 7.2 percent. Thus, in these data, all beetle effect and response were both highly correlated and showed phylogenetic signal (λ = 0.67).

**Figure 1.**
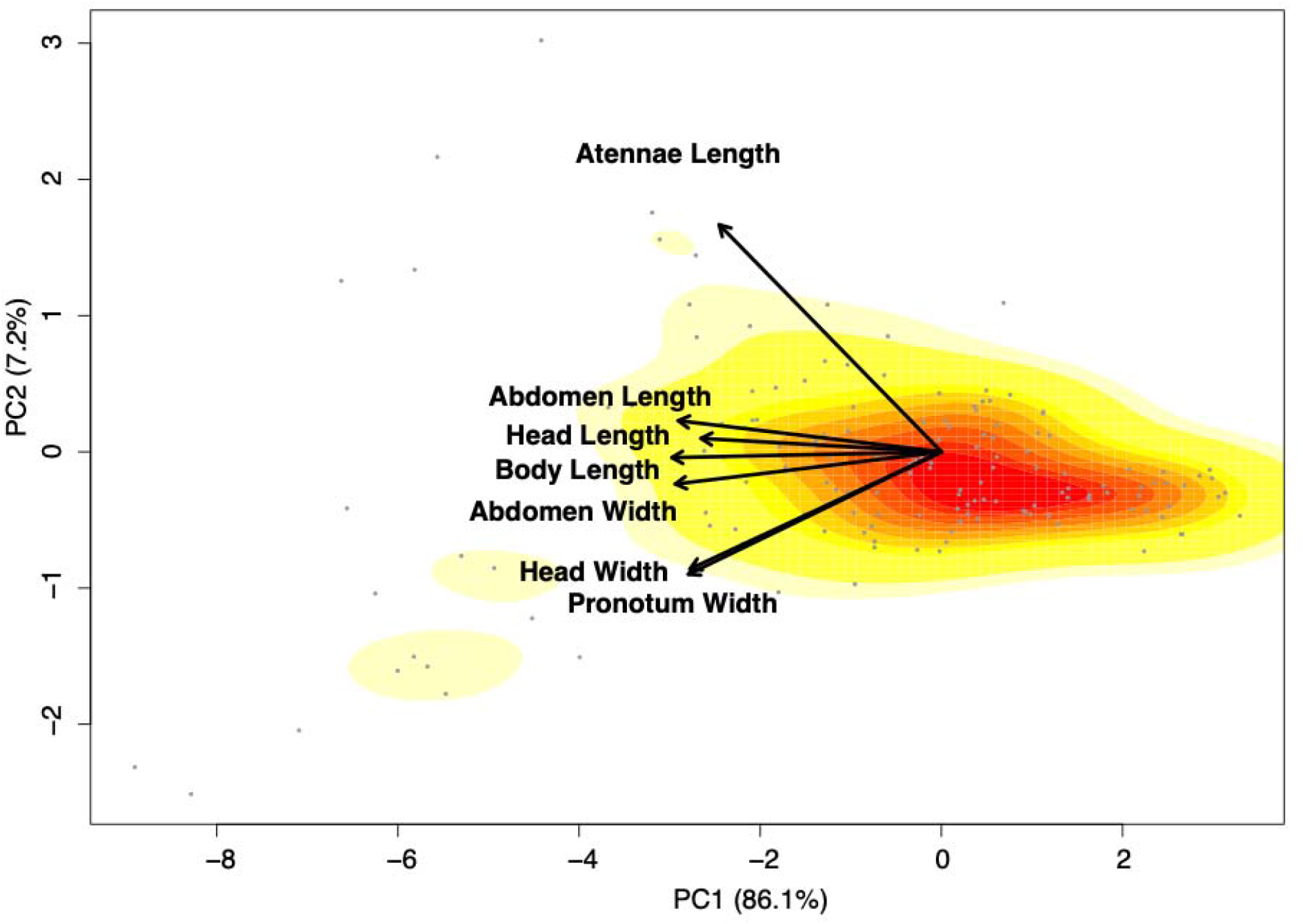
Phylogenetic PCA representing seven morphological ground beetle traits. Points reflect each of 154 species, and arrows represent the loadings of the morphological traits. Background color shading indicates the density of points within the space. PC1 represents 86.1% of the total variance among traits, and PC2 represents an additional 7.2% (full summary and loadings in Appendix S7). Head and body length, and abdomen length and width all load strongly onto PC1, while PC2 captures additional variation in pronotum and head width and antennae length. There is a strong correlation among all traits, including effect traits (head length, body length) and response traits (all traits).

**Table 2.** Measure of phylogenetic signal (Pagel’s λ, for all traits p < 0.001), and AIC weights for seven evolutionary models, for each of seven morphological ground beetle traits. Models include Brownian motion (BM), Ornstein-Uhlenbeck (OU), early burst (EB), trend through time (TR), Pagel’s λ, Pagel’s δ, and white noise (WN). In all traits, Pagel’s λ values were > 0.73 and Pagel’s λ was the most-likely model *(i.e.,* had the lowest AIC; δAIC > 56.1). Strong support for Pagel’s λ is consistent with trait evolution intermediate between pure chance and pure Brownian motion, *i.e.*, phylogenetic signal.

| Trait | Pagel's $\lambda$ | BM | OU | EB | TR | $\lambda$ | $\delta$ | WN |
| --- | --- | --- | --- | --- | --- | --- | --- | --- |
| Body Length | 0.82 | 1204.0 | 967.0 | 1206.0 | 1160.4 | 892.4 | 1127.2 | 970.4 |
| Abdomen Length | 0.86 | 1015.5 | 815.5 | 1017.5 | 974.0 | 742.2 | 943.4 | 819.7 |
| Abdomen Width | 0.81 | 939.4 | 689.1 | 941.4 | 895.0 | 617.5 | 860.9 | 694.0 |
| Pronotum Width | 0.76 | 952.2 | 659.0 | 954.2 | 906.0 | 578.4 | 869.8 | 661.4 |
| Head Length | 0.75 | 647.8 | 375.6 | 649.8 | 602.9 | 317.9 | 568.1 | 377.6 |
| Head Width | 0.74 | 785.0 | 534.2 | 787.0 | 739.7 | 453.3 | 704.5 | 536.7 |
| Antennae Length | 0.88 | 775.3 | 670.3 | 777.3 | 742.6 | 614.1 | 721.6 | 683.3 |

## Discussion

We found evidence of phylogenetic signal in all ground beetle traits measured, and our AIC-based theoretic model comparison best supported Pagel’s λ for all seven traits. These results indicate that these North American ground beetles’ morphologies are evolutionarily conserved, and phylogenetic inertia is a stronger driver of trait evolution than lineages’ recent or independent responses to ecological disturbances and interactions (Cooper et al. 2010). Furthermore, phylogenetic PCA revealed that all measured body shape traits, both within and across the effect and response categories, are highly correlated. The λ value of the PCA reinforced our finding of strong phylogenetic signal in individual beetle traits, consistent (again) with conserved trait evolution along the principal component axes. Together, these results pose a conservation concern: the correlation of beetle effect and response traits suggests that a severe disturbance event may lead to a decline in ecosystem services (Díaz et al. 2013), and shared traits among closely-related species increases the risk of disturbance leading to the loss of entire clades.

### Evolutionary constraints on ground beetle morphology

The strong phylogenetic signal (λ > 0.73) we identified across seven morphological ground beetle traits is consistent with the findings of previous studies, which have found evidence of phylogenetic signal in traits for mammals (Arnaudo et al. 2019), trees (Swenson et al. 2017) and even habitat-specialized beetle clades (Vamosi and Vamosi 2007). In these other studies, and ours, links between phylogenetic structure in traits and key ecological processes suggest that traits were evolved early in species’ evolutionary history and helped define clades’ relationships with their environments. Researchers have also found evidence that beetle assemblages are phylogenetically clustered (e.g., Funk et al. 1995), and that predator-prey relationships in ground beetle assemblages are conserved (Brousseau et al. 2018). This further highlights the importance of considering phylogenetic constraint in predictions of how environmental change might lead to the loss of beetle diversity and associated ecological functions.

We note that, while our study has broad spatial coverage (much of the United States of America), our taxonomic sampling is biased. Pitfall traps were used to collect these specimens; our beetles are ecologically similar and thus a non-random subset of all North American species. Other studies attempting to apply phylogenetic risk-assessment are likely to encounter similar biases, and so we urge caution in uncritically applying our results, and those of similarly biased samples, to broader macro-evolutionary contexts. That said, such studies represent valuable pieces in the macro-evolutionary puzzle, and our study contributes new clade-specific information to our growing knowledge of beetle evolution as a whole. Testing the taxonomic and geographic scale-dependence of phylogenetic signal body traits across beetle families is an important next step, but requires a complete phylogeny and is therefore outside of the scope of the present study.

### Correlation among functional traits

The effect-response trait framework provides an opportunity to investigate ecological responses to environmental change using functional traits as a proxy for how organisms affect and are affected by their environments (Suding et al. 2008). Phylogenetic signal in effect or response traits, as we found in this study, may represent an increased risk that environmental stressors will lead to the loss of multiple closely-related beetles from an ecosystem (Díaz et al. 2013). In addition to the risk associated with conserved evolution of these traits, the strong correlations among traits demonstrated by our phylogenetic PCA suggests further ecosystem risk. When effect and response traits are linked, as in our results, the loss of traits associated with environmental change is more likely to correspond with the loss of traits linked to particular ecological functions. Despite the novelty of applying the effect-response trait framework to beetles (Fountain-Jones et al. 2015) and in particular, the need for more experimental research to investigate effect traits (Wong et al. 2019), the correlation among all traits measured suggests ecological response and function are broadly linked. Seemingly small changes in a species’ environment could therefore lead to changes in community structure and function (Mayer and Rietkerk 2004, Groffman et al. 2006). For North American beetles, this loss of function could result in increased harmful pathogen presence, aphid predation on crops, fire-interval variability, loss of biodiversity, or a loss of key habitat health indicators (Logan and Powell 2001, Rainio and Niemelä 2003). Understanding evolutionary constraints on and correlations among other ecologically important traits, such as those associated with beetle behavior and adaptive responses to environmental change, are important topics for future research.

For the many ecosystems reliant on services provided by ground beetles, this effect-response framework provides a useful insight for conservation practitioners: saving species will matter. Those species that are most likely to be lost are providing different ecosystem services from those that are left behind, and so preserving natural beetle assemblages is worthwhile if beetle ecosystem services are a priority.

In conclusion, we find evidence of strong phylogenetic signal in the North American ground beetle traits we measured, suggesting the evolution of these traits is constrained by phylogenetic inertia. While our dataset is expansive (covering 154 species), it does not reflect all Carabidae (or even all of those found within North America). These results provide a unique contribution to our understanding of evolutionary conservatism across phylogenetic scales, as well as the past and future of ecological assembly in this particular clade. The strong correlation among effect and response traits in this study implies that local extinction events could lead to a loss of ecosystem services. Resource managers who want to preserve ecosystem function in the face of global change should therefore attempt to preserve beetle assemblages as a whole. We hope that the data we release with this manuscript will be of use in future studies examining drivers of beetle co-occurrence, invasion and dispersal success, and the controls on beetle range limits. We call for further work to examine whether species’ effect and response traits can be distinguished among beetle taxa, particularly in studies examining a more ecologically and taxonomically broad set of species.

## Data Accessibility Statement

All data and code used in this study are included in the supplementary materials.

## Acknowledgments

We thank the National Ecological Observatory Network, particularly Katie LeVan and Kate Thibault, for data sharing and support. AK, ASG, HW, JDS, WDP, and the Pearse Lab are supported by National Science Foundation grants ABI-1759965 and EF-1802605, and UKRI/NERC NE/V009710/1.

## Supplementary Materials

**Appendix S1.** R script used for all statistical analyses and figures presented in the manuscript.

**Appendix S2.** Collection locations for ground beetle specimens collected and photographed by the National Ecological Observatory Network (NEON). For each of 4886 specimens, this database includes a unique NEON ID, sample ID, taxonomic classification, information about the identification and vouchering of the specimen, demographic information (life stage and sex), and details pertaining to the site of collection, including habitat, latitude, longitude, elevation, and NEON domain. To further explore the map of NEON sites, see: https://www.neonscience.org/field-sites/field-sites-map

**Appendix S3.** Morphological trait measurements for 1975 ground beetle (Carabidae) specimens. Traits were measured from photographs (one per specimen) taken by the National Ecological Observatory Network (NEON). The traits included are body length, abdomen length, abdomen width, pronotum width, head length, head width, and antennae length. All measurements were recorded in millimeters and represent the maximum possible length or width of the trait.

**Appendix S4.** Protocol developed and used in this study for measuring seven morphological traits from dorsal images of beetle specimens.

**Appendix S5.** Cytochrome oxidase one (COX1) sequences for 2411 specimens of ground beetles collected by NEON. This database includes a NEON ID, taxonomic classification, and COX1 sequence for each of the specimens digitized and measured in this study (and additional specimens).

**Appendix S6.** 1000 bootstrap phylogenies of 2399 beetle sequences and five outgroup sequences (from *Myrmeleon immaculatus, M. formicarius, Brachynemurus abdominalis, B. ferox,* and *Dendroleon obsoletus),* used to conduct independent maximum likelihood searches using RAxML.

**Appendix S7.** Phylogeny of 2399 beetle sequences and five outgroup sequences (from *Myrmeleon immaculatus, M. formicarius, Brachynemurus abdominalis, B. ferox,* and *Dendroleon obsoletus*). Sequences were sampled by NEON and dated as described in the text.

**Appendix S8.** Summary and loadings of phylogenetic PCA for seven ground beetle functional traits. Effect traits are noted with an asterisk (all of the effect traits in this study also function as response traits).

## Literature Cited

Albert, J., M. Platek, and L Cizek. 2012. Vertical stratification and microhabitat selection by the Great Capricorn Beetle *(Cerambyx cerdo)* (Coleoptera: Cerambycidae) in open-grown, veteran oaks. European Journal of Entomology 109:553–559.

Arnaudo, M. E., N. Toledo, L. Soibelzon, and P. Bona. 2019. Phylogenetic signal analysis in the basicranium of Ursidae (Carnivora, Mammalia). PeerJ 7:e6597.

Barton, P. S., H. Gibb, A. D. Manning, D.B. Lindenmayer, and S. A. Cunningham. 2011. Morphological traits as predictors of diet and microhabitat use in a diverse beetle assemblage. Biological Journal of the Linnean Society 102:301–310.

Bentz, B. J., J. Régnière, C. J. Fettig, E. M. Hansen, J. L. Hayes, J. A. Hicke, R. G. Kelsey, J. F. Negrón, S. J. Seybold. 2010. Climate Change and Bark Beetles of the Western United States and Canada: Direct and Indirect Effects. BioScience 60:602–613.

Boettiger, C., G. Coop, and P. Ralph. 2012. Is Your Phylogeny Informative? Measuring the Power of Comparative Methods. Evolution 66:2240–2251.

Brousseau, P.-M., D. Gravel, and I. T. Handa. 2018. Trait matching and phylogeny as predictors of predator–prey interactions involving ground beetles. Functional Ecology 32:192–202.

Butler, M. A., and A. A. King. 2004. Phylogenetic Comparative Analysis: A Modeling Approach for Adaptive Evolution. The American Naturalist 164:683–695.

Cardoso, P., T. L. Erwin, P. A. V. Borges, and T. R. New. 2011. The seven impediments in invertebrate conservation and how to overcome them. Biological Conservation 144:2647–2655.

Cavender-Bares, J., K. H. Kozak, and S. W. Kembel. 2009. The merging of community ecology and phylogenetic biology. Ecology Letters 12:693–715.

Collins, K. L., N. D. Boatman, A. Wilcox, J. M. Holland, and K. Chaney. 2002. Influence of beetle banks on cereal aphid predation in winter wheat. Agriculture, Ecosystems & Environment 93:337–350.

Cooper, N., W. Jetz, and R. P. Freckleton. 2010. Phylogenetic comparative approaches for studying niche conservatism. Journal of Evolutionary Biology 23:2529–2539.

Díaz, S., A. Purvis, J. H. C. Cornelissen, G. M. Mace, M. J. Donoghue, R. M. Ewers, P. Jordano, W. D. Pearse. 2013. Functional traits, the phylogeny of function, and ecosystem service vulnerability. Ecology and Evolution 3:2958–2975.

Eastman, J. M., M. E. Alfaro, P. Joyce, A. L. Hipp, and L. J. Harmon. 2011. A Novel Comparative Method for Identifying Shifts in the Rate of Character Evolution on Trees. Evolution 65:3578–3589.

Elgar, M.A., D. Zhang, Q. Wang, B. Wittwer, H. T. Pham, T. L. Johnson, C. B. Freelance, and M. Coquilleau. 2018. Focus: ecology and evolution: insect antennal morphology: the evolution of diverse solutions to odorant perception. The Yale Journal of Biology and Medicine, 91:457.

Fountain Jones, N. M., S. C. Baker, and G. J. Jordan. 2015. Moving beyond the guild concept: developing a practical functional trait framework for terrestrial beetles. Ecological Entomology 40:1–13.

Fountain Jones, N. M., G. J. Jordan, C. P. Burridge, T. J. Wardlaw, T. P. Baker, L. Forster, M. Petersfield, S. C. Baker. 2017. Trophic position determines functional and phylogenetic recovery after disturbance within a community. Functional Ecology 31:1441–1451.

Funk, D. J., D. J. Futuyma, G. Ortí, and A. Meyer. 1995. Mitochondrial DNA sequences and multiple data sets: a phylogenetic study of phytophagous beetles (Chrysomelidae: Ophraella). Molecular Biology and Evolution 12:627–640.

Gossner, M. M., T. Lachat, J. Brunet, G. Isacsson, C. Bouget, H. Brustel, R. Brandl, W. Weisser, J. Mueller. 2013. Current Near-to-Nature Forest Management Effects on Functional Trait Composition of Saproxylic Beetles in Beech Forests. Conservation Biology 27:605–614.

Groffman, P. M., J. S. Baron, T. Blett, A. J. Gold, I. Goodman, L. H. Gunderson, B. M. Levinson, M. A. Palmer, H. W. Paerl, G. D. Peterson, N. L. Poff, D. W. Rejeski, J. F. Reynolds, M. G. Turner, K. C. Weathers, and J. Wiens. 2006. Ecological Thresholds: The Key to Successful Environmental Management or an Important Concept with No Practical Application? Ecosystems 9:1–13.

Haila, Y., I. K. Hanski, J. Niemelä, P. Punttila, S. Raivio, and H. Tukia. 1994. Forestry and the boreal fauna: matching management with natural forest dynamics. Annales Zoologici Fennici 31:187–202.

Hance, T., C. Grégoire-Wibo, and P. Lebrun. 1990. Agriculture and ground-beetles populations. The consequence of crop types and surrounding habitats on activity and species composition. Pedobiologia 34:337–346.

Harmon, L. J., J. A. Schulte, A. Larson, and J. B. Losos. 2003. Tempo and Mode of Evolutionary Radiation in Iguanian Lizards. Science 301:961–964.

Harmon, L. J., J. T. Weir, C. D. Brock, R. E. Glor, and W. Challenger. 2008. GEIGER: investigating evolutionary radiations. Bioinformatics 24:129–131.

Harmon, L. J., J. B. Losos, T. J. Davies, R. G. Gillespie, J. L. Gittleman, W. B. Jennings, K. H. Kozak, M. A. McPeek, F. Moreno-Roark, T. J. Near, A. Purvis, R. E. Ricklefs, D. Schluter, J. A. Schulte II, O. Seehausen, B. L. Sidlauskas, O. Torres-Carvajal, J. T. Weir, A. Ø. Mooers. 2010. Early Bursts of Body Size and Shape Evolution Are Rare in Comparative Data. Evolution 64:2385–2396.

Hebert, P. D. N., S. Ratnasingham, and J. R. de Waard. 2003. Barcoding animal life: cytochrome c oxidase subunit 1 divergences among closely related species. Proceedings of the Royal Society of London. Series B: Biological Sciences 270:S96–S99.

Hortal, J., J. A. F. Diniz-Filho, L. M. Bini, M. Á. Rodríguez, A. Baselga, D. Nogués-Bravo, T. F. Rangel, B. A. Hawkins, and J. M. Lobo. 2011. Ice age climate, evolutionary constraints and diversity patterns of European dung beetles: Ice age determines European scarab diversity. Ecology Letters 14:741–748.

Hunt, T., J. Bergsten, Z. Levkanicova, A. Papadopoulou, O. S. John, R. Wild, P. M. Hammond, D. Ahrens, M. Balke, M. S. Caterino, J. Gómez-Zurita, I. Ribera, T. G. Barraclough, M. Bocakova, L. Bocak, A. P. Vogler. 2007. A Comprehensive Phylogeny of Beetles Reveals the Evolutionary Origins of a Superradiation. Science 318:1913–1916.

Ives, A. R., and H. C. J. Godfray. 2006. Phylogenetic Analysis of Trophic Associations. The American Naturalist, 168:E1–E14.

Katoh, K. 2002. MAFFT: a novel method for rapid multiple sequence alignment based on fast Fourier transform. Nucleic Acids Research 30:3059–3066.

Kotze, D. J., P. Brandmayr, A. Casale, E. Dauffy-Richard, W. Dekoninck, M. J. Koivula, G. L. Lövei, D. Mossakowski, J. Noordijk, W. Paarmann, R. Pizzolotto, P. Saska, A. Schwerk, J. Serrano, J. Szyszko, A. Taboada, H. Turin, S. Venn, R. Vermeulen, T. Zetto. 2011. Forty years of carabid beetle research in Europe – from taxonomy, biology, ecology and population studies to bioindication, habitat assessment and conservation. ZooKeys 100:55–148.

Logan, J. A., and J. A. Powell. 2001. Ghost Forests, Global Warming, and the Mountain Pine Beetle (Coleoptera: Scolytidae). American Entomologist 47:160–173.

Luck, G. W., S. Lavorel, S. McIntyre, and L. Lumb. 2012. Improving the application of vertebrate trait-based frameworks to the study of ecosystem services. Journal of Animal Ecology 81:1065–1076.

Mayer, A. L., and M. Rietkerk. 2004. The Dynamic Regime Concept for Ecosystem Management and Restoration. BioScience 54:1013–1020.

McGill, B., B. Enquist, E. Weiher, and M. Westoby. 2006. Rebuilding community ecology from functional traits. Trends in Ecology & Evolution 21:178–185.

Moretti, M., A. T. C. Dias, F. de Bello, F. Altermatt, S. L. Chown, F. M. Azcárate, J. R. Bell, B. Fournier, J. Hortal, S. Ibanez, Öckinger E, J. P. Sousa, J. Ellers, M. P. Berg. 2017. Handbook of protocols for standardized measurement of terrestrial invertebrate functional traits. Functional Ecology 31:558–567.

Münkemüller, T., S. Lavergne, B. Bzeznik, S. Dray, T. Jombart, K. Schiffers, and W. Thuiller. 2012. How to measure and test phylogenetic signal. Methods in Ecology and Evolution 3:743–756.

Muschick, M., A. Indermaur, and W. Salzburger. 2012. Convergent Evolution within an Adaptive Radiation of Cichlid Fishes. Current Biology 22:2362–2368.

Pagel, M. 1999. Inferring the historical patterns of biological evolution. Nature 401:877–884.

Poff, N. L., J. D. Olden, N. K. M. Vieira, D. S. Finn, M. P. Simmons, and B. C. Kondratieff. 2006. Functional trait niches of North American lotic insects: traits-based ecological applications in light of phylogenetic relationships. Journal of the North American Benthological Society 25:730–755.

Rainio, J., and J. Niemelä. 2003. Ground beetles (Coleoptera: Carabidae) as bioindicators. Biodiversity & Conservation 12:487–506.

Revell, L. J., L. J. Harmon, and D. C. Collar. 2008. Phylogenetic signal, evolutionary process, and rate. Systematic Biology 57:591–601.

Revell, L. J. 2009. Size-Correction and Principal Components for Interspecific Comparative Studies. Evolution 63:3258–3268.

Ribera, I., S. Dolédec, I. S. Downie, and G. N. Foster. 2001. Effect of Land Disturbance and Stress on Species Traits of Ground Beetle Assemblages. Ecology 82:1112–1129.

Savage, V. M., C. T. Webb, and J. Norberg. 2007. A general multi-trait-based framework for studying the effects of biodiversity on ecosystem functioning. Journal of Theoretical Biology 247:213–229.

Schindelin, J., I. Arganda-Carreras, E. Frise, V. Kaynig, M. Longair, T. Pietzsch, S. Preibisch, C. Rueden, S. Saalfeld, B. Schmid, J.-Y. Tinevez, D. J. White, V. Hartenstein, K. Eliceiri, P. Tomancak, A. Cardona. 2012. Fiji: an open-source platform for biological-image analysis. Nature Methods 9:676–682.

Schmitz, O. 2017. Predator and prey functional traits: understanding the adaptive machinery driving predator–prey interactions. F1000Research 6:1767.

Seehausen, O. 2006. African cichlid fish: a model system in adaptive radiation research. Proceedings of the Royal Society B: Biological Sciences 273:1987–1998.

Smith, S. A., and B. C. O’Meara. 2012. treePL: divergence time estimation using penalized likelihood for large phylogenies. Bioinformatics 28:2689–2690.

Spake, R., N. Barsoum, A. C. Newton, and C. P. Doncaster. 2016. Drivers of the composition and diversity of carabid functional traits in UK coniferous plantations. Forest Ecology and Management 359:300–308.

Stamatakis, A. 2014. RAxML version 8: a tool for phylogenetic analysis and post-analysis of large phylogenies. Bioinformatics 30:1312–1313.

Suding, K. N., S. Lavorel, F. S. Chapin, J. H. C. Cornelissen, S. Díaz, E. Garnier, D. Goldberg, D. U. Hooper, S. T. Jackson, and M.-L. Navas. 2008. Scaling environmental change through the community-level: a trait-based response-and-effect framework for plants. Global Change Biology 14:1125–1140.

Swenson, N. G., M. D. Weiser, L. Mao, M. B. Araújo, J. A. F. Diniz□Filho, J. Kollmann, D. Nogués-Bravo, S. Normand, M. A. Rodríguez, R. García-Valdés, F. Valladares, M. A. Zavala, and J.-C. Svenning. 2017. Phylogeny and the prediction of tree functional diversity across novel continental settings. Global Ecology and Biogeography 26:553–562.

Tucker, C. M., M. W. Cadotte, S. B. Carvalho, T. J. Davies, S. Ferrier, S. A. Fritz, R. Grenyer, M. R. Helmus, L. S. Jin, A. O. Mooers, S. Pavoine, O. Purschke, D. W. Redding, D. F. Rosauer, M. Winter, F. Mazel. 2017. A guide to phylogenetic metrics for conservation, community ecology and macroecology. Biological Reviews 92:698–715.

Uyeda, J. C., & L. J. Harmon. 2014. A novel Bayesian method for inferring and interpreting the dynamics of adaptive landscapes from phylogenetic comparative data. Systematic Biology 63:902–918.

Vamosi, J. C., & S. M. Vamosi. 2007. Body size, rarity, and phylogenetic community structure: insights from diving beetle assemblages of Alberta. Diversity and Distributions 13:1–10.

Vandewalle, M., F. De Bello, M. P. Berg, T. Bolger, S. Doledec, F. Dubs, C. K. Feld, R. Harrington, P. A. Harrison, S. Lavorel, and P. M. Da Silva. 2010. Functional traits as indicators of biodiversity response to land use changes across ecosystems and organisms. Biodiversity and Conservation 19:2921–2947.

Violle, C., M.-L. Navas, D. Vile, E. Kazakou, C. Fortunel, I. Hummel, and E. Garnier. 2007. Let the concept of trait be functional! Oikos 116:882–892.

Webb, C. O., D. D. Ackerly, M. A. McPeek, and M. J. Donoghue. 2002. Phylogenies and Community Ecology. Annual Review of Ecology and Systematics 33:475–505.

Wiens, J. J., D. D. Ackerly, A. P. Allen, B. L. Anacker, L. B. Buckley, H. V. Cornell, E. I. Damschen, T. J. Davies, J. A. Grytnes, S. P. Harrison, B. A. Hawkins, R. D. Holt, C. M. McCain, P. R. Stephens. 2010. Niche conservatism as an emerging principle in ecology and conservation biology. Ecology Letters 13:1310–1324.

Wong, M. K. L., B. Guénard, and O. T. Lewis. 2019. Trait-based ecology of terrestrial arthropods. Biological Reviews 94:999–1022.

Zhang, S.-Q., L.-H. Che, Y. Li, D. Liang, H. Pang, A. Ślipiński, and P. Zhang. 2018. Evolutionary history of Coleoptera revealed by extensive sampling of genes and species. Nature Communications 9:1–11.

