## Supplementary material for "Strong trait correlation and phylogenetic signal in North American ground beetle (Carabidae) morphology": Appendix_S1.R

```
# Code for Stachewicz et al., 2020. "Strong trait correlation and phylogenetic 
# signal in North American ground beetle (Carabidae) morphology"

################################
# Libraries ####################
################################
library(ks)        # For densities in PCA plots
library(caper)     # Keeping tree and data aligned
library(phytools)  # Fitting lambda/K and geologic ages plotting
library(geiger)    # Model fitting

################################
# Loading data #################
################################
# Load trait data and match names (excluding those missing specific IDs)
image.lookup <- with(read.csv("Appendix_S2_SpecimenData.csv", as.is=TRUE),
               setNames(species_name, processid))
data <- read.csv("Appendix_S3_TraitData.csv", as.is=TRUE)
data$Species <- image.lookup[data$ImageID]
data <- data[data$Species != "",]
data <- data[,c("Species","Trait","Value")]

# Calculate species' median values
data <- with(data, tapply(Value, list(Species, Trait), median))
data <- as.data.frame(data)
data$Species <- rownames(data)

# Load phylogeny and match its names
entire.phylo <- phylo <- read.tree("Appendix_S6_Phylogeny.tre")
phylo$tip.label <- gsub(".", "-", phylo$tip.label, fixed=TRUE)
phylo.lookup <- with(read.csv("Appendix_S5_SequenceMetadata.csv", as.is=TRUE),
                     setNames(name, gene))
names(phylo.lookup) <- unlist(substr(names(phylo.lookup), 0, 10))
phylo$tip.label <- unname(phylo.lookup[phylo$tip.label])
phylo <- drop.tip(phylo, which(is.na(phylo$tip.label))) # Remove the outgroups
phylo.for.figure <- phylo
phylo <- drop.tip(phylo, which(duplicated(phylo$tip.label))) # Remove duplicates of species
phylo <- force.ultrametric(phylo) # Correct numerical precision in branch-lengths

# Make comparative dataset to match traits and phylogeny
c.data <- comparative.data(data =data, phy = phylo, names.col = Species)

################################
# Univariate analysis ##########
################################
# Estimate signal in all traits
sapply(c.data$data, function(x) phylosig(c.data$phy, x, method="lambda", test=TRUE))
sapply(c.data$data, function(x) phylosig(c.data$phy, x, method="K", test=TRUE))

# Estimate best-fit model in all traits
.aics.models <- function(trait, tree){
    modes <- c("BM","OU","EB","trend","lambda","delta","drift","white")
    models <- lapply(modes, function(x) fitContinuous(tree, setNames(trait,tree$tip.label), model=x))
    return(setNames(sapply(models, function(x) x$opt$aic), modes))
}
model.aics <- sapply(c.data$data, .aics.models, tree=c.data$phy)

################################
# (Phylo) PCA ##################
################################
# Run phylogenetic PCA (note: scaling makes little difference)
names(c.data$data) <- c("Atennae Length", "Abdomen Length", "Abdomen Width", 
                        "Body Length", "Head Length", "Head Width", "Pronotum Width")
p.pca <- phyl.pca(c.data$phy, scale(c.data$data), method="lambda")
p.pca$lambda
summary(p.pca)
biplot(p.pca)

# Run regular PCA
pca <- prcomp(c.data$data, scale=TRUE)

# Plot the output from the Phylogenetic PCA
.pca.plot <- function(pca, dims, scale=5, ...){
    if(inherits(pca, "phyl.pca")){
        scores <- pca$S
        rotation <- pca$L
        traits <- rownames(rotation)
    } else {
        scores <- pca$x
        rotation <- pca$rotation
        traits <- rownames(rotation)
    }
    density <- kde(scores[,dims])
    plot(scores[,dims], type="n", axes=FALSE)
    axis(1)
    axis(2)
    plot(density, display="filled.contour", cont=seq(10,90,10), cols=heat.colors(length(seq(10,90,10))), add=TRUE, axes=FALSE)
    points(scores, pch=20, cex=.5, col="grey60")
    arrows(rep(0,ncol(scores)),rep(0,ncol(scores)), rotation[,dims[1]]*scale, rotation[,dims[2]]*scale, length=.1, lwd=3)
    text.locs <- rotation*scale*1.3
    text(text.locs[,dims[1]], text.locs[,dims[2]], traits, font=2)
}

.pca.plot(p.pca, 1:2, scale=3)

################################
# Phylogeny figure for MS ######
################################
plotTree(phylo.for.figure, ftype="off",ylim=c(-0.1*Ntip(phylo.for.figure),Ntip(phylo.for.figure)), lwd=.75)
geo.legend()
```
