## Supplementary material for "Strong trait correlation and phylogenetic signal in North American ground beetle (Carabidae) morphology": Appendix_S4_Protocol.docx

**FIJI Beetle Trait Measurement Protocol**

*All measurements are taken from a dorsal view*

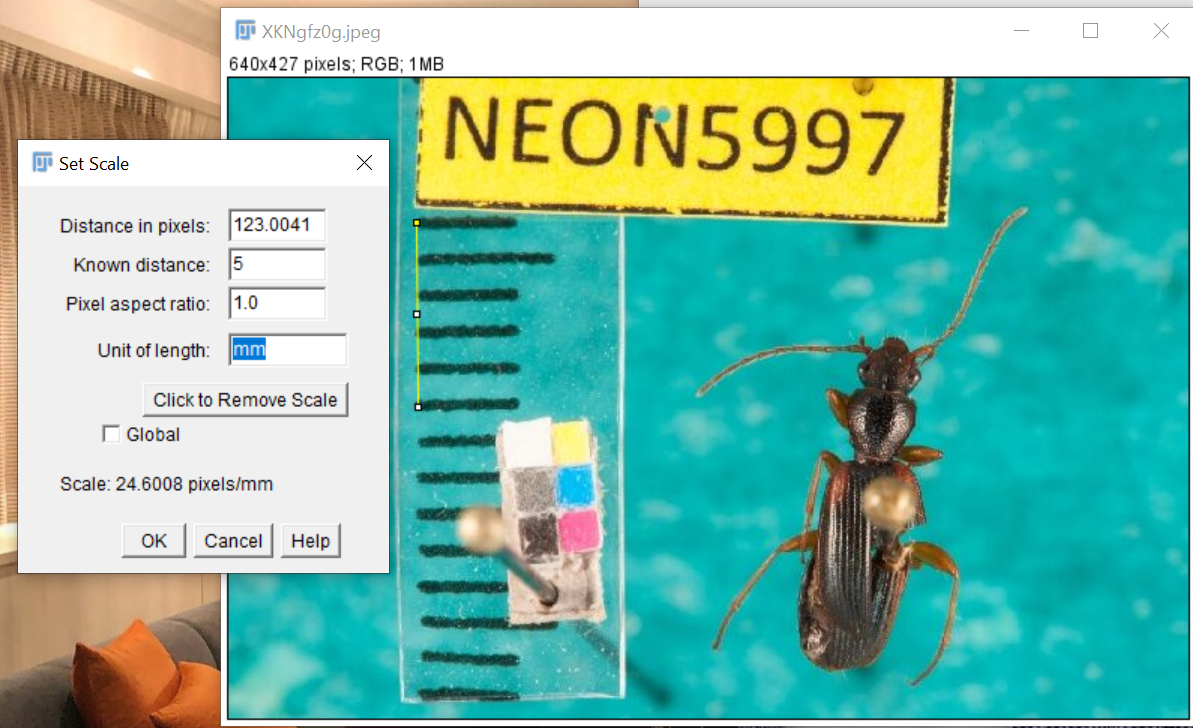
**Set 5mm scale**: Utilizing the scale ruler within the image, use the straight-line tool and from the most bottom edge (to reduce angle error) count out 5 mm, then set the known distance using the set scale option.

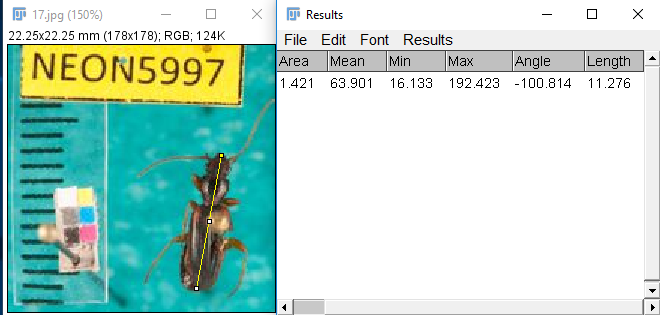
**Maximum Body Length:** Using the straight-line tool draw from the tip of the abdomen to the end of the clypeus.

**Maximum Abdomen Length**: Use the straight-line tool to draw a line from the wing tip of the abdomen to scutellum.

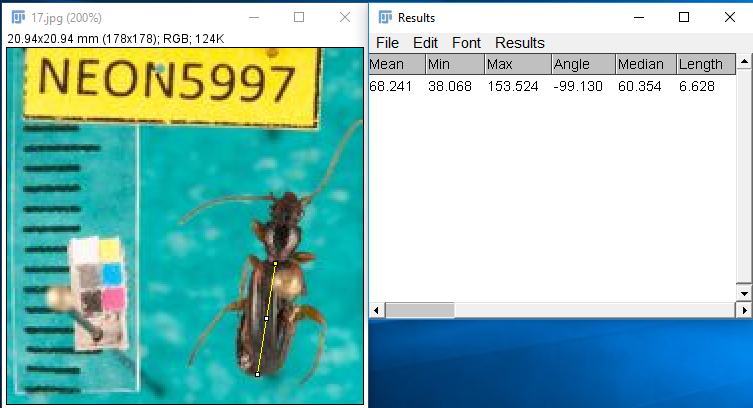

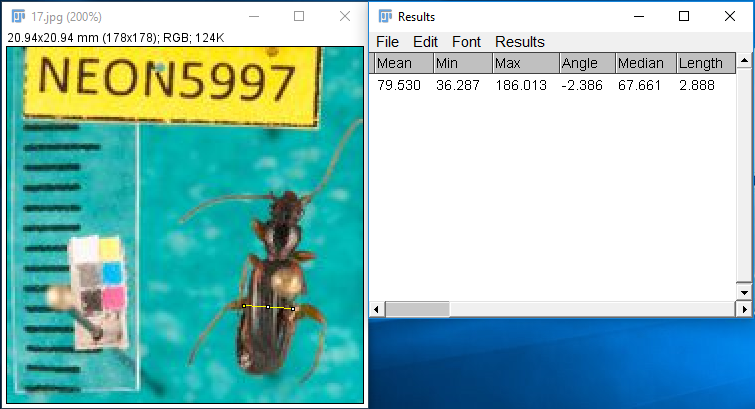
**Maximum Abdomen Width**: Determine the widest area of the abdomen and measure across using the straight-line tool.

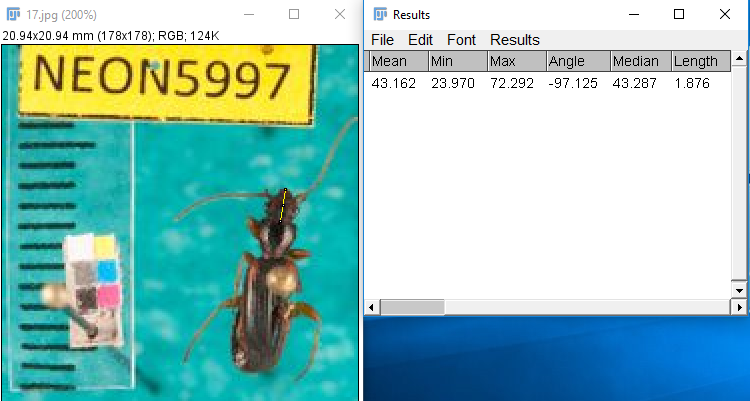
**Maximum Head Length**: Use the straight-line tool to draw a line down the middle of beetle head from the head-pronotum joint to just before the mandible if mandible is visible.

**Maximum Head Width**: Identify the widest point of the head and use the straight-line tool to measure across.

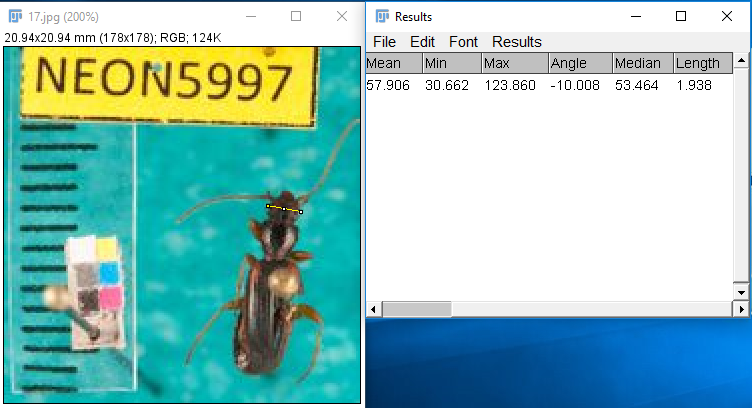

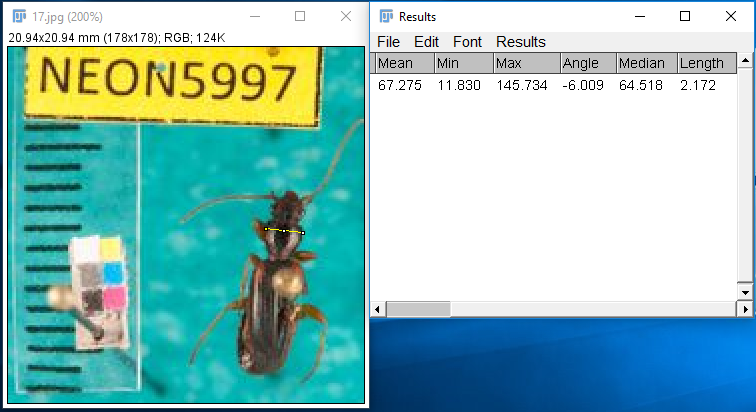
**Maximum Pronotum Width**: Identify the widest area of the pronotum and measure straight across using the straight-line tool.

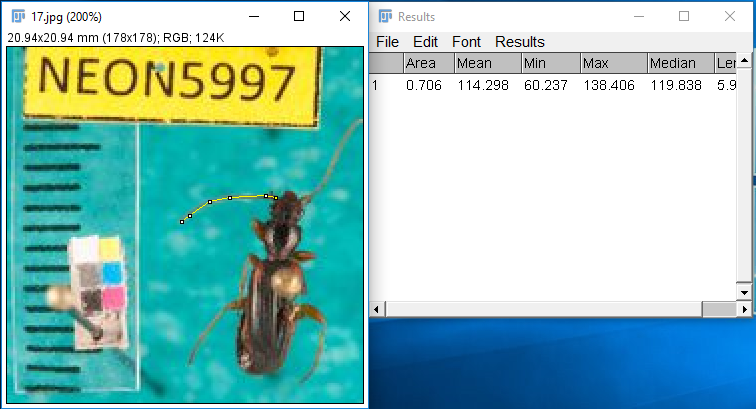
**Antennae Length**: Use the segmented line tool and trace entire antennae from antennae tip to scape.
