## Supplementary material for "Strong trait correlation and phylogenetic signal in North American ground beetle (Carabidae) morphology": Supplement_S8_PCA.docx

**Appendix S7**

Table 1. Importance of components in Phylogenetic PCA of seven morphological ground beetle traits.

|  | PC1 | PC2 | PC3 | PC4 | PC5 | PC6 | PC7 |
| --- | --- | --- | --- | --- | --- | --- | --- |
| Standard Deviation | 0.151 | 0.044 | 0.033 | 0.019 | 0.016 | 0.008 | 0.004 |
| Proportion of Variance | 0.861 | 0.072 | 0.040 | 0.013 | 0.009 | 0.003 | 0.001 |
| Cumulative Proportion | 0.861 | 0.933 | 0.974 | 0.987 | 0.997 | 0.999 | 1.000 |

Table 2. Loadings for Phylogenetic PCA of seven morphological ground beetle traits. Loadings are presented for the first three components as they represent a cumulative 97.4% of the variation among all traits. Effect traits are noted with an asterisk (all of the effect traits in this study also function as response traits).

| Trait | PC1 (86.1%) | PC2 (7.2%) | PC3 (4.0%) |
| --- | --- | --- | --- |
| Antennae Length* | -0.818 | 0.556 | -0.076 |
| Abdomen Length | -0.968 | 0.076 | -0.136 |
| Abdomen Width | -0.979 | -0.079 | -0.108 |
| Body Length | -0.992 | -0.014 | -0.044 |
| Head Length* | -0.884 | 0.033 | 0.464 |
| Head Width | -0.925 | -0.287 | 0.006 |
| Pronotum Width* | -0.931 | -0.300 | -0.115 |
